# Impaired islet function with normal exocrine enzyme secretion is consistent across the head, body, and tail pancreas regions in type 1 diabetes

**DOI:** 10.1101/2024.02.08.579175

**Authors:** Denise M. Drotar, Ana Karen Mojica-Avila, Drew T. Bloss, Christian M. Cohrs, Cameron T. Manson, Amanda L. Posgai, MacKenzie D. Williams, Maigan A. Brusko, Edward A. Phelps, Clive H. Wasserfall, Stephan Speier, Mark A. Atkinson

## Abstract

Histopathological heterogeneity in human pancreas has been well documented; however, functional evidence at the tissue level is scarce. Herein we investigated *in situ* glucose-stimulated islet and carbachol-stimulated acinar cell secretion across the pancreas head (PH), body (PB), and tail (PT) regions in no diabetes (ND, n=15), single islet autoantibody-positive (1AAb+, n=7), and type 1 diabetes donors (T1D, <14 months duration, n=5). Insulin, glucagon, pancreatic amylase, lipase, and trypsinogen secretion along with 3D tissue morphometrical features were comparable across the regions in ND. In T1D, insulin secretion and beta-cell volume were significantly reduced within all regions, while glucagon and enzymes were unaltered. Beta-cell volume was lower despite normal insulin secretion in 1AAb+, resulting in increased volume-adjusted insulin secretion versus ND. Islet and acinar cell secretion in 1AAb+ were consistent across PH, PB and PT. This study supports low inter-regional variation in pancreas slice function and potentially, increased metabolic demand in 1AAb+.

## Introduction

In type 1 diabetes (T1D), islet infiltrating immune cells cause destruction of the pancreatic beta-cells in a patchy, lobular pattern^1,2^. Recent efforts have demonstrated that other islet cell types (e.g., alpha-cells^3–5^) and pancreatic compartments (e.g., acini^6–8^) may also be impacted during T1D pathogenesis. Indeed, aberrant glucagon levels were described in individuals with T1D^9–11^, and alpha-cell failure in response to hypoglycemia was proposed as a possible mechanism^3^. Moreover, exocrine changes have been noted during the development of T1D, where the pancreas weight/volume is decreased in individuals with T1D, islet autoantibody-positive individuals without diabetes (AAb+), and healthy first-degree relatives of T1D patients^12–16^. Moreover, serum trypsinogen and lipase levels are also significantly decreased in pre-T1D (marked by the presence of ≥2AAb) as well as in recent-onset and established T1D individuals^17,18^. Whether this is due to reduced number of acinar cells^7^, individual acinar cell size^6^, or an intrinsic functional defect is not clear.

Extensive characterization of pancreata from individuals with no diabetes (ND) has unveiled differences in islet size, cellular composition, and density across the three main regions of the organ^19–22^. Specifically, large islets contain increased proportion of alpha-cells, whereas small islets are almost exclusively comprised of beta-cells^19^. Alpha and beta-cell frequencies are reduced in the pancreas head (PH) compared to body (PB) and tail (PT) due to increased numbers of pancreatic polypeptide-secreting cells, which are usually restricted to the uncinate process within the PH^21^. Islet density is similar within the PH and PB, yet significantly increased in the PT^22^. The functional relevance of these morphological differences across the organ is still unclear, and whether pancreatic heterogeneity plays a role in the development of diseases such as T1D remains to be investigated.

Pancreas tissue function is difficult to study, especially in the human setting, and most of our knowledge is based on systemic measurements of serological markers in response to standardized glucose or mixed meal challenges. Understanding of human islet biology and pathophysiology has increased dramatically in recent decades with improved access to human pancreas tissues via the Network for Pancreatic Organ donors with Diabetes (nPOD)^23,24^, Human Pancreas Analysis Program (HPAP), and Integrated Islet Distribution Program (IIDP)^25^. Isolated islet preparations from ND donors are widely used to assess islet function, yet generally lack information regarding islet localization or the local microenvironment, as isolations are typically performed from the whole organ rather than regions, with a few exceptions such as the DiViD studies on laparoscopically procured pancreas tail biopsies^26–28^. Moreover, islet isolation from T1D pancreata is challenging and prone to bias towards structurally intact islets that survive the isolation process^28^. Alternative approaches, such as the live tissue slice preparation, now enable investigation of islets within their tissue environment from T1D, AAb+, and ND donors^29–32^. Slices from pancreatic tissue retain both the endocrine and exocrine compartments, with islets surrounded by acinar cells^33^, preserved vasculature^31,34^, and extracellular matrix (ECM) components^35^. This preparation facilitates the investigation of the collective function of endocrine cells irrespective of the disease stage (i.e., immune infiltration and degree of beta-cell destruction) as well as acinar tissue from the same donor or sample. Endocrine-exocrine crosstalk is increasingly recognized as a key facet of diabetes pathogenesis as well as other diseases of the pancreas^36^. Indeed, the interplay between these compartments is crucial for proper organ development and growth, and it is reasonable to posit that a deficit in one of these will directly affect its counterpart^37^.

We hypothesized that the reported differences in islet cell density and composition among the three pancreas regions^20,21^ might influence islet and acinar cell secretion patterns or capacity, or impact the pathogenesis of diseases such as diabetes. To test this, we performed an extensive functional study of the three pancreatic regions in humans by investigating insulin and glucagon release, as well as pancreatic enzymes secreted by the acinar tissue from the same donors using live pancreas tissue slices. We characterized islet function and volume, as well as acinar cell function averaged across 8-12 slices per region from the PH, PB, and PT of 15 ND donors, 7 single AAb+ [1AAb+; including glutamic acid decarboxylase AAb+ (GADA+, n=6) and insulin AAb+ (IAA+, n=1)], and 5 recent-onset T1D donors with disease duration <14 months (donor characteristics in Table S1 and Fig. S1A-D).

## Results

### Similar hormone secretion, islet composition, and enzyme secretion across PH, PB and PT in human ND pancreas

Hormone release was assessed in freshly prepared tissue slices from the PH, PB and PT of 15 ND donors. Glucose stimulation of live pancreas slices elicited the expected two-phase insulin secretion kinetics in all three regions of the pancreas at both medium (5.5 mM) and high glucose concentrations (11.1 mM), referred to as 5.5G and 11.1G, respectively, as well as following membrane depolarization (30 mM KCl) (Fig. 1A). Although no significant differences were identified across regions, we noted inter-donor variability across the PH, PB and PT.

**Fig 1.**
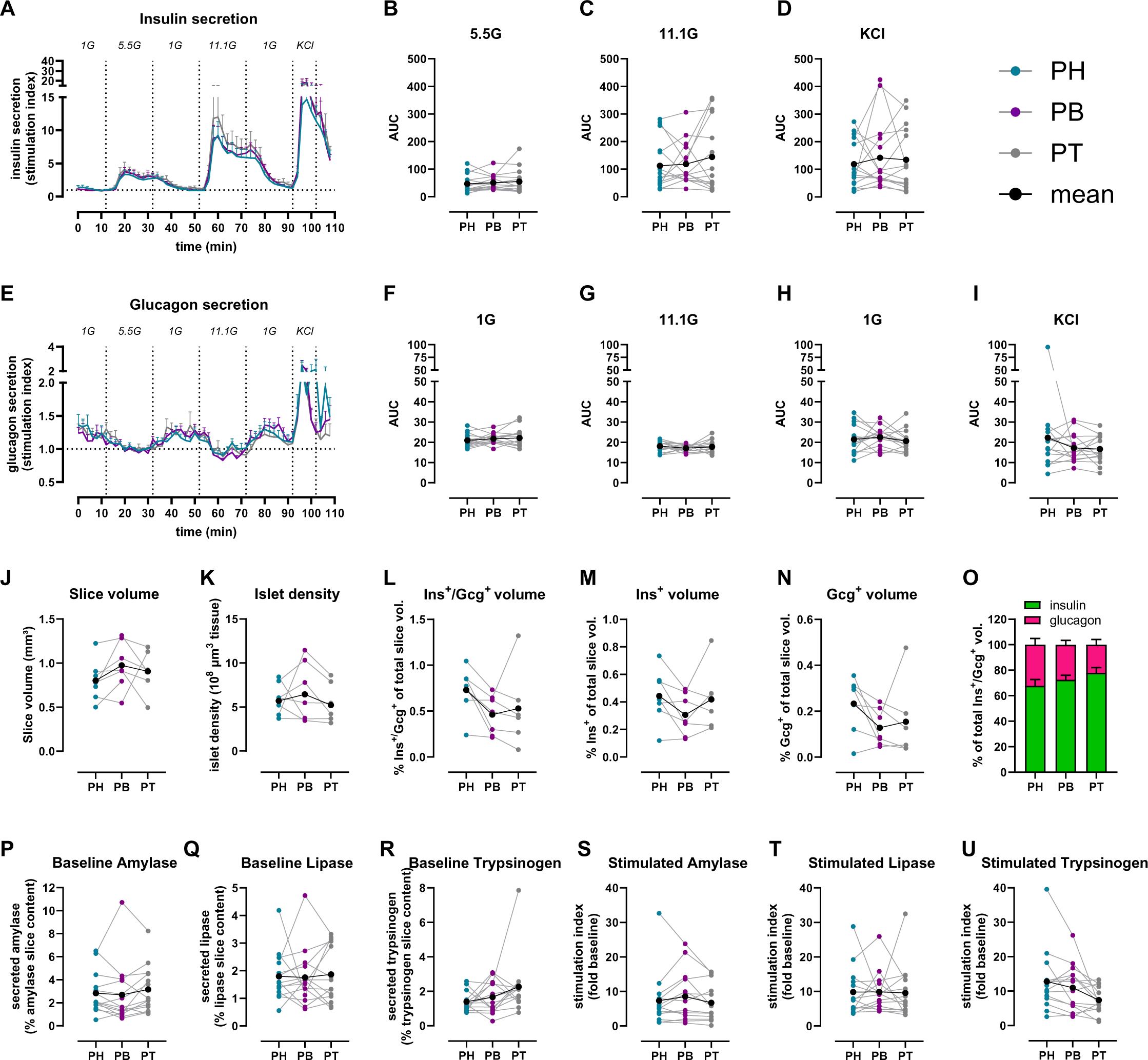
Similar islet and acinar cell secretion across the PH, PB and PT in the human ND pancreas. A & E. Insulin (A) and glucagon (E) secretion traces from slices of PH (teal), PB (purple), and PT (grey) shown as stimulation index calculated as fold of mean baseline (1G for insulin and 5.5G for glucagon). B-D. Quantification of insulin responses to 5.5G (B), 11.1G (C), and KCl (D) stimulation. F-I. Quantification of glucagon responses to 1G (F), 11.1G (G), 1G (H), KCl (I) stimulation. J-O. 3D morphometry of perifused slices showing slice volume (J), islet density (K), endocrine (L), insulin (M), and glucagon (N) volumes of perifused slices, and the contribution of insulin and glucagon volume to the total endocrine volume in perifused slices (O). P-R. Baseline pancreatic amylase (P), lipase (Q), and trypsinogen (R) release expressed as % of total enzyme content. S-U. Stimulated amylase (S), lipase (T), and trypsinogen (U) secretion in response to 10 µM carbachol expressed as stimulation index calculated as fold of baseline. N=15 ND donors for A-I, 7 ND donors for J-O, with 4 slices/region/donor and 14 ND donors for P-U with 5 slices/region/donor Dots represent individual donors, means are shown in black. Data are represented as mean ± SEM. Significance was assessed using RM one-way ANOVA of log-transformed data (B-D, F-I, J-N and P-U). ∗p < 0.05; ∗∗p < 0.01; ∗∗∗p < 0.001; ∗∗∗∗p < 0.0001; ns, not significant.

Quantifying insulin responsiveness to all applied stimuli (glucose or KCl) showed similar secretion output in slices prepared from the different regions of the ND pancreas (Fig. 1B-D). Glucagon secretion was measured at the same timepoints as insulin, and normalized to the 5.5G step in order to investigate both stimulatory and suppressive effects of glucose (Fig. 1E). Glucagon release in response to decreases in glucose concentration (5.5G to 1G, or 11.1G to 1G) and KCl were comparable across PH, PB and PT (Fig. 1F, H-I) with suppression of glucagon release upon high glucose infusion also showing no differences across the regions in slices from ND donors (Fig. 1G).

Next, we employed tissue 3D morphometric analysis across whole tissue slices after functional assessment to quantify total insulin and glucagon volumes from islets, clusters and single cells in 7 ND pancreata. Overall, slice tissue volumes used for perifusion and 3D morphometric quantification were comparable across the three regions from ND donors (Fig. 1J). Our whole slice 3D morphometric analysis resulted in similar islet density across the PH, PB and PT in ND (Fig. 1K), as well as comparable endocrine volume in these slices, regardless of the region from which the tissue was obtained (Fig. 1L-N). Insulin contributed 70-80% to the observed endocrine volume in all ND pancreas regions (Fig. 1O).

We also investigated acinar cell secretory function using adjacent slices obtained from the same donors and regions, comparing pancreatic amylase, lipase and trypsinogen release. Under basal conditions, similar secretion of all three enzymes was observed across the PH, PB and PT in ND donors (Fig. 1P-R). We then examined enzyme release upon stimulation with 10 µM carbachol, a widely used stimulus of acinar cell secretion^33^. Although secretion of all pancreatic enzymes increased ∼10-fold in the presence of carbachol, stimulated enzyme levels did not significantly differ across regions (Fig. 1S-U). Altogether, these data indicate that in *ex vivo* pancreas tissue slices, islets and acinar cells have similar secretion responses, independent of the region of origin within the ND pancreas.

### Diminished insulin, normal glucagon secretion across all pancreatic regions in recent-onset T1D

We next investigated whether T1D might differentially affect insulin and glucagon secretion within the PH, PB, and PT regions. Insulin responses in slices from the 1AAb+ group were similar to ND in all pancreas regions (Fig. 2A-I, Fig. S2A-E, orange lines, and Fig. S3A-E). In comparison to both control and 1AAb+ donor groups, glucose-stimulated insulin secretion was dramatically diminished in all 5 recent-onset T1D cases investigated, with loss of the typical two-phase insulin response^3,29^ (Fig. 2A-I and Fig. S2A-E, red lines). Specifically, two donors had undetectable insulin secretion, particularly from the PH and PB (Fig. S3F-J), but overall, there were no significant differences in insulin responsiveness across the regions in T1D slices. Furthermore, upon normalization to the baseline, insulin responses to 5.5G were completely lost in slices from all pancreas regions. Interestingly, 2 out of 5 T1D cases still showed weak 11.1G peaks (Fig. 2D-H, and Fig. S3I). Moreover, some slices from 3 recent-onset T1D donors exhibited a near-normal response to KCl (Fig. 2D-F, I), but the responsive region varied for each case (Fig. S3J). Altogether, these data suggest reduced glucose sensitivity and preserved insulin secretion capacity in these donors.

**Fig 2.**
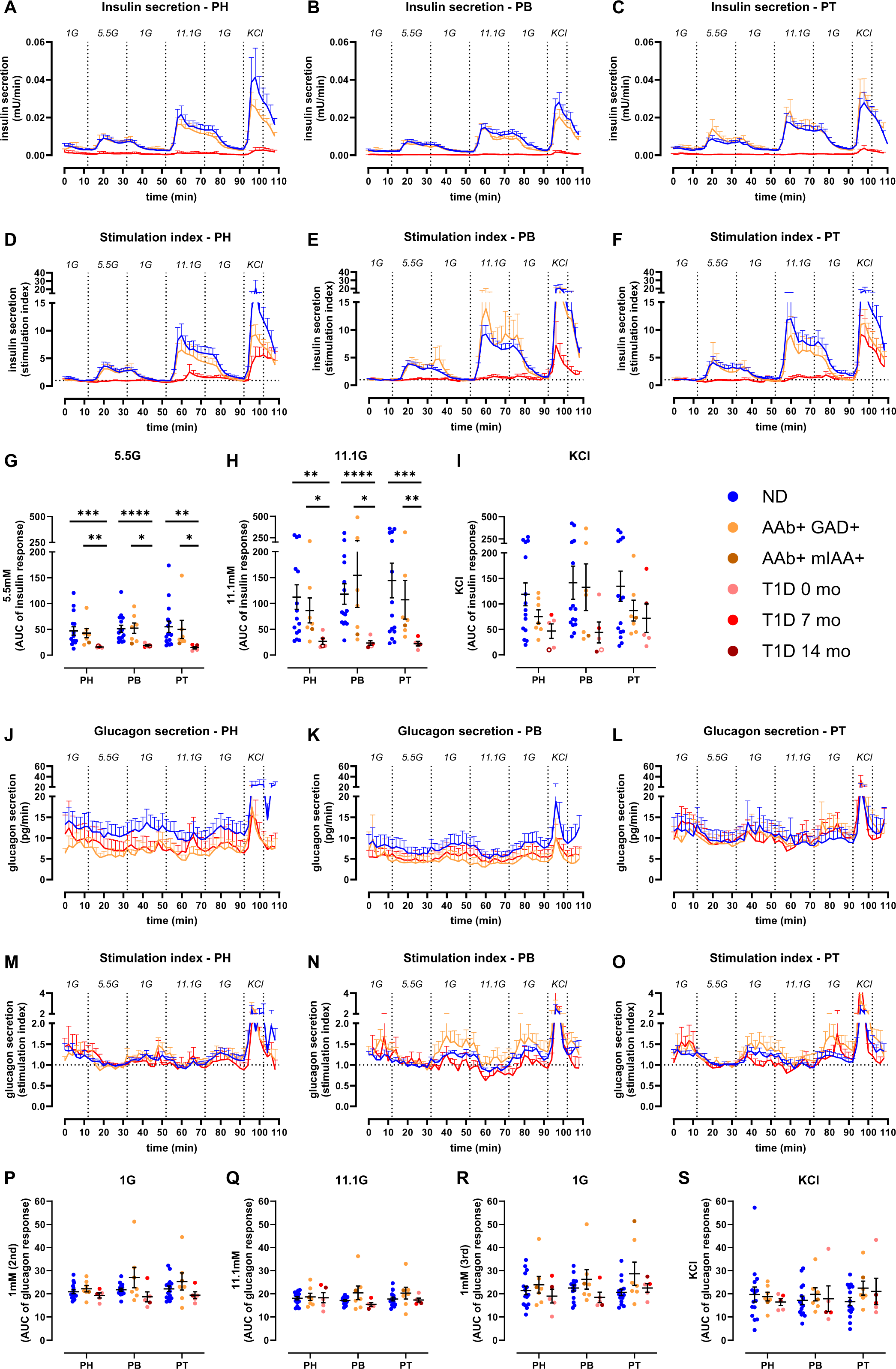
Reduced insulin but not glucagon secretion in all pancreas regions in recent-onset T1D. A-C. Insulin secretion from slices of PH (A), PB (B), and PT (C) in slices from ND, 1AAb+, and T1D donors shown as absolute secreted amounts (mU/min). D-F. Insulin secretion traces from slices of PH (D), PB (E), and PT (F) in slices from ND, 1AAb+, and T1D donors shown as stimulation index calculated as fold of baseline (1G). G-I. Quantification of insulin responses to 5.5G (G), 11.1G (H), and KCl (I) stimulation. J-L. Glucagon secretion from slices of PH (J), PB (K), and PT (L) in slices from ND, 1AAb+, and T1D donors shown as absolute secreted amounts (pg/min). M-O. Glucagon secretion traces from slices of PH (M), PB (N), and PT (O) in slices from ND, 1AAb+, and T1D donors shown as stimulation index calculated as fold of baseline (5.5G). P-S. Quantification of glucagon responses to 1G (P), 11.1G (Q), 1G (R), and KCl (S) stimulation. N=15 ND, 7 1AAb+, and 5 T1D donors, with 4 slices/region/donor Dots represent individual donors. Data are represented as mean ± SEM. Significance was assessed using RM two-way ANOVA of log-transformed data (G-I and P-S). ∗p < 0.05; ∗∗p < 0.01; ∗∗∗p < 0.001; ∗∗∗∗p < 0.0001; ns, not significant. See also Figures S2 and S3.

Glucagon secretion was measured in the same slices from these T1D, 1AAb+, and ND donors, represented in Fig. 2J-O. Absolute amounts of glucagon release were variable (Fig. 2J-L); however, after normalization to the 5.5G baseline, glucagon responses to changes in glucose concentrations did not differ significantly across disease groups (Fig. 2M-S and Fig. S2F-K) or pancreatic regions (Fig. S3K-V).

### Preserved acinar cell function at T1D onset

Using the same approach as noted above for ND donors, we measured pancreatic enzyme release in slices from each region of 1AAb+ and recent-onset T1D donors. Basal release of pancreatic amylase, lipase, and trypsinogen was comparable in slices from 1AAb+, T1D, and ND pancreata, across all regions (Fig. 3A-C). Similarly, following carbachol stimulation, enzyme secretion increased ∼10-fold across all regions in the 1AAb+ and T1D groups (Fig. 3D-F), and the stimulation index was neither substantially different amongst the disease groups, nor the pancreatic regions (Fig. 3G-I).

**Fig 3.**
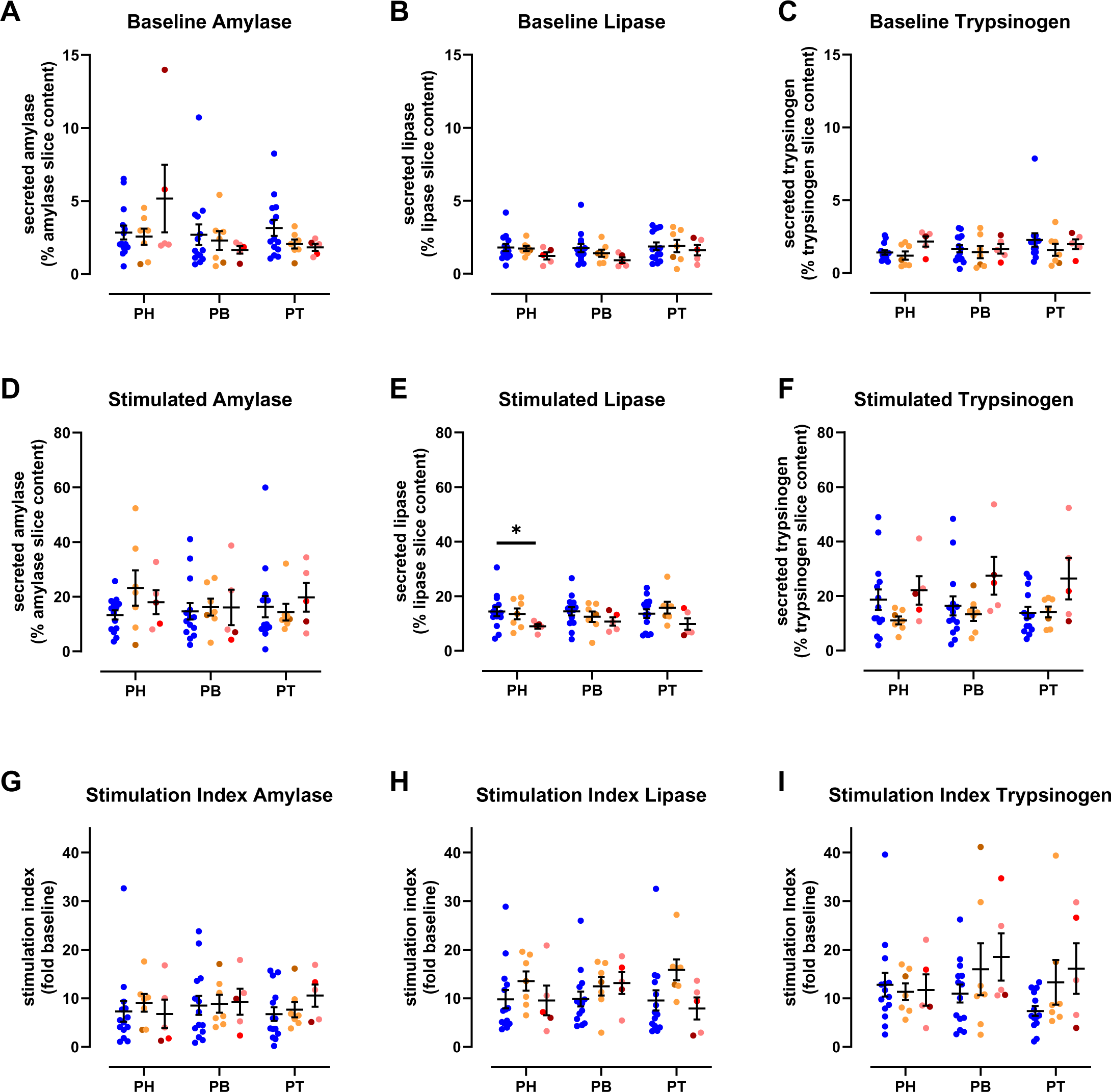
Preserved acinar cell function at T1D onset. A-C. Baseline pancreatic amylase (A), lipase (B), and trypsinogen (C) release in ND, 1AAb+, and T1D slices from the PH, PB, and PT, expressed as % of total enzyme content. D-F. Stimulated amylase (D), lipase (E), and trypsinogen (F) secretion in response to carbachol in ND, 1AAb+, and T1D slices from PH, PB, and PT, expressed as % of total enzyme content. G-I. Stimulated amylase (G), lipase (H), and trypsinogen (I) secretion in response to 10 µM carbachol in ND, 1AAb+, and T1D slices from PH, PB, and PT, expressed as stimulation index calculated as fold of baseline. N=14 ND, 7 1AAb+, and 5 T1D donors, with 5 slices/region/donor Dots represent individual donors. Data are represented as mean ± SEM. Significance was assessed using RM two-way ANOVA. ∗p < 0.05; ∗∗p < 0.01; ∗∗∗p < 0.001; ∗∗∗∗p < 0.0001; ns, not significant.

### Decreased endocrine and beta-cell volume across the pancreas at T1D onset

In order to better understand the relationship between mass and function of the endocrine and exocrine pancreas compartments at the onset of T1D, we performed 3D morphometric analysis of the perifused slices. Slices prepared from PH, PB and PT of 7 ND, 5 1AAb+, and 5 recent-onset T1D donors had similar total tissue slice volumes (Fig. 4A). The endocrine cell volume (sum of insulin and glucagon) was significantly decreased in both 1AAb+ and T1D cases (Fig. 4B) due to a significant reduction of beta-cell volume together with normal alpha-cell volume in PH and PB (Fig. 4C-D). Islet density appeared to be lower in 3 T1D cases but did not reach statistical significance (p= 0.2290 for PH, p=0.3115 for PB and p=0.3666 for PT as compared to ND) (Fig. 4E). While total endocrine cell volume was not significantly reduced in the PT as noted above, our analysis shows a reduced beta to alpha-cell ratio in slices from the PT of T1D donors (Fig. 4F-H).

**Fig 4.**
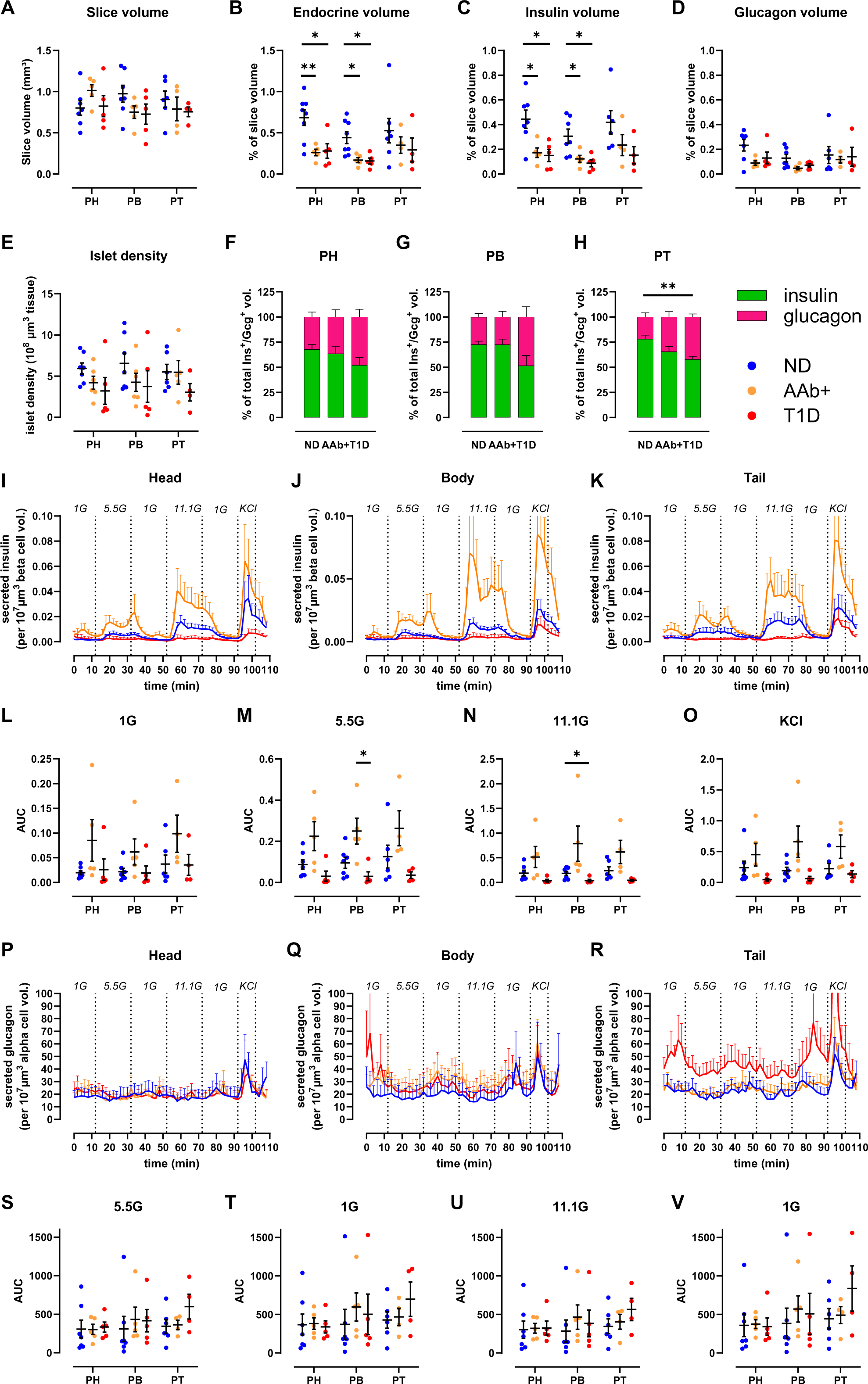
Decreased endocrine and beta-cell mass across the pancreas at T1D onset. A-H. 3D morphometry of perifused slices showing slice volume (A), endocrine (B), insulin (C), and glucagon (D) volumes of perifused slices, islet density (E), and the contribution of insulin and glucagon volume to the total endocrine volume across the disease groups in PH (F), PB (G), and PT (H). I-K. Insulin secretion traces from slices of PH (I), PB (J), and PT (K) in slices from ND, 1AAb+, and T1D donors shown as secreted insulin normalized to beta-cell volume in the respective slices. L-O. Quantification of insulin responses to 1G (L), 5.5G (M), 11.1G (N), and KCl (O) stimulation. P-R. Glucagon secretion traces from slices of PH (P), PB (Q), and PT (R) in slices from ND, 1AAb+, and T1D donors shown as secreted glucagon normalized to alpha-cell volume in the respective slices. S-V. Quantification of glucagon responses to 5.5G (S), 1G (T), 11.1G (U), and 1G (V) stimulation. N=7 ND, 5 1AAb+, and 5 T1D donors, with 4 slices/region/donor Dots represent individual donors. Data are represented as mean ± SEM. Significance was assessed using RM two-way ANOVA of log-transformed data. ∗p < 0.05; ∗∗p < 0.01; ∗∗∗p < 0.001; ∗∗∗∗p < 0.0001; ns, not significant.

Next, we used the volumetric data to normalize hormone secretion in tissue slices where both parameters were measured. This approach allows the investigation of insulin secretion of the remaining beta-cell volume within the slices (Fig. 4I-K). In all pancreas regions, beta-cell volume-normalized insulin secretion was diminished in slices from recent-onset T1D, particularly in response to high glucose stimulation (Fig 4L-O). In contrast, because beta-cell volume was decreased in 1AAb+ slices (Fig. 4C) while overall secretion was preserved (Fig. 2A-I), the beta-cell volume-adjusted insulin secretion appeared to be amplified in some 1AAb+ slices when compared to ND (Fig 4I-K). Importantly, none of the 1AAb+ cases investigated herein showed signs of dysglycemia, as serum HbA1c and C-peptide levels were normal (Fig. S1). We also used alpha-cell volumes to normalize glucagon secretion within the same slices. When adjusted to the glucagon positive alpha-cell volume, glucagon secretion was consistent across all regions in ND and 1AAb+ (Fig. 4P-R). PT slices of recent-onset T1D showed slightly increased alpha-cell volume-normalized glucagon release (Fig. 4R), but did not reach significance in our analysis (Fig. 4S-V).

## Discussion

Here we have performed an extensive functional study on both islet and acinar cells across the three human pancreas regions in ND, 1AAb+, and recent-onset T1D donors. We demonstrated that glucose stimulation produced a typical two-phase insulin release pattern, as previously described^29,38^, in each pancreas region separately, as well as when regions were taken together. To date, we are aware of only one study that compared glucose-stimulated insulin release in isolated islets from the PH + PB versus the PT^20^. Our results from living tissue slices confirm that insulin release is similar across the pancreatic regions in ND, and expand into glucagon release as well as pancreatic enzyme secretion, providing first-time investigation of both endocrine and exocrine secretion within the same tissues.

Interestingly, despite known developmental, morphological, and histological differences across the main pancreatic regions (e.g., percentage or proportion of beta, alpha, delta or PP-cells)^20–22^, the cases we investigated herein demonstrated similar hormone secretion from islets and pancreatic enzyme release from acini across the PH, PB and PT of ND donors. Indeed, we did not observe differences in endocrine cell mass or composition, nor in islet cell density in slices across the regions. These results differ from previously published extensive histomorphometric studies on ND human pancreata where islet density was increased in the PT when compared to PH or PB, and the frequency of alpha and beta-cells was diminished in PH^20,21^. These disparities are likely due to differences in sample preparation and size (thin whole organ cross-sections versus tissue slices of 120 µm thickness), analysis approach (2D versus 3D in this study), and human donor variability.

We extended these regional studies on pancreata obtained from 1AAb+ and recent-onset T1D under the hypothesis that during the pathogenesis of T1D, some regions might exhibit a greater degree of dysfunction than others; this, based on the well-recognized phenomenon of inconsistent and lobular patterns of immune infiltration and destruction of islets during T1D progression^2^. While limited to 1AAb+ cases (six GADA+ and one IAA+) due to donor availability, this is the first study investigating islet and acinar cell function in slices from AAb+ donors, who are considered to have increased T1D risk^39,40^. Both absolute and baseline normalized insulin secretion from 1AAb+ slices were similar to ND in PH, PB, and PT. 3D morphometric analysis of the perifused slices allowed us to investigate endocrine cell volume within the same tissue pieces. As insulin-positive beta-cell volume was significantly decreased in slices from PH and PB of 1AAb+ donors, the mass-adjusted insulin release was rather increased in this group, in all regions, suggesting that these beta-cells are perhaps experiencing a higher workload to maintain glycemic control. Interestingly, it was previously shown that some 1AAb+ individuals present signs of beta-cell dysfunction years before T1D clinical diagnosis^41^, but whether or when our 1AAb+ donors would have developed T1D is, of course, unknown. Data from additional at-risk donors are needed, particularly those with two or more islet AAb, in order to gain insight into the mechanisms that lead to the loss of beta-cell function during T1D progression.

In slices from recent-onset T1D donors (<14 months disease duration), we observed a dramatic loss of insulin release in response to glucose across the PH, PB, and PT. Interestingly, responses to KCl depolarization were preserved in 3 of these cases, suggesting a defect in glucose responsiveness rather than insulin production, confirming our previous observations^29^. This decrease in insulin release can be explained by the significant reduction in insulin-positive beta-cell volume in the PH and PB (Fig. 4C). Although this was not statistically significant in the PT, we did find significant changes in the alpha and beta-cell mass proportions in T1D versus ND slices. Nevertheless, beta-cell volume-normalized insulin release was dramatically reduced in all regions for most T1D donors, suggesting that the remaining beta-cell mass is functionally impaired. This finding corroborates previous studies using slices from T1D donors^29^, but differs from other studies using isolated human islets where residual beta-cells retained features of regulated insulin secretion^4,5^. These observations could be driven by differences in experimental tissue preparation (e.g., isolated islets versus tissue slices, including mechanical and chemical stressors during isolation, recovery culture times or potential selectivity bias towards intact islets), stimuli (amino-acids or IBMX versus glucose), and analysis approach (normalization to islet numbers/islet equivalent (IEQ) versus baseline or morphometry).

In addition to investigating insulin secretion across all pancreas regions, we also examined glucagon release within the same samples. We did not find differences in glucagon secretion or alpha-cell volumes across the PH, PB and PT of ND donors, nor did we identify differences across the ND, 1AAb+, and T1D donor groups suggesting that alpha-cell volume and function are preserved in these 1AAb+ and recent-onset T1D cases. This contrasts with previous studies that reported dysregulated glucagon secretion from alpha-cells within tissue slices^3^ and isolated human islets^4,5^. Besides differences in experimental approach described above (tissue preparation, stimuli or analysis and normalization approaches), perhaps the most noteworthy difference is donor age and diabetes duration. Specifically, the studies referenced above investigated islet functionality in cases with established T1D (up to 30 years duration) while we focused on recent onset T1D, with most of our donors diagnosed at demise.

The contribution of the exocrine compartment in T1D has gained interest over the past decade, sparked initially by observations of decreased pancreas size, not only in persons with long-standing T1D but also in AAb+ and first-degree relatives of individuals with T1D^12–14,16^. At the functional level, it has been shown that circulating levels of pancreatic amylase, lipase, and trypsinogen^17,18^ and fecal levels of elastase-1^42,43^ are significantly reduced even before clinical diagnosis, but whether this is resultant of a functional deficit at the acinar cell level is not fully understood^44^. Leveraging unique tissue access, we performed an *in situ* investigation of acinar cell secretion in living slices from our cohort of ND, 1AAb+, and recent-onset T1D donors. We measured both baseline and carbachol-stimulated secretion of pancreatic enzymes from the PH, PB, and PT and found no significant difference across groups or regions. These results suggest that the acinar cell secretory function might not be affected and rather, point to reduced pancreas organ mass— which is well-documented in established, recent-onset, and pre- T1D^12,13,16^— and reduced acinar cell numbers^7^ as the likely reason for lower pancreatic enzyme levels in serum and stool during T1D progression^17,18^.

Finally, while our study is the first to investigate islet and acinar cell secretion across the main pancreatic regions in ND, 1AAb+, and T1D donors, we are aware this poses certain limitations. In this study, live tissue slice preparation was restricted to a specific area within each region (i.e., one cross-section per region), therefore it might not fully reflect overall organ biology or pathophysiology. To overcome this, we prepared approximately 40 ± 15 slices from several tissue blocks within each cross-section, and 10 slices were randomly selected for functional assays to provide an impartial representation of that region. We were also limited in the number of donor pancreata available for research; therefore, exact case matching (for age, sex, BMI, HLA) was not possible. For this reason, we strived to conduct these studies on every donor organ received by nPOD within the past three years, and only excluded cases that had an exceptional medical history and could not be categorized into the standard groups of ND, AAb+ or T1D by nPOD (e.g., endocrine tumors, MODY).

Taken together, our data demonstrate that islet and acinar cell secretion profiles were comparable between PH, PB, and PT in ND. Insulin secretion was dramatically impaired in recent-onset T1D subjects, which was consistent across the pancreas regions. Alpha-cell and acinar cell secretion were unaltered in 1AAb+ and T1D donors, independent of pancreas region. Further studies are needed on pancreas slices from rare pre-T1D donors, defined as having two or more AAb, in order to characterize the progressive loss of islet cell function across the disease stages. Although no striking inter-regional disparities were observed and despite the shortcomings associated with human tissue research, we performed the most extensive studies to date of both islet and acinar cell function in the three pancreas regions in ND human donors. Moreover, these regional studies were extended to extremely rare pancreata from individuals at the very early phases of T1D pathogenesis. Finally, we believe that this study contributes to a better understanding of the endocrine and exocrine pancreas function in the human setting and possibly the mechanisms that lead to the lobular heterogeneity of insulitis and beta-cell loss in organ donors with T1D.

## Supporting information

Key Resources Table

## Acknowledgements

We would like to thank the nPOD Administrative and Organ Processing and Pathology Core (OPPC) teams, respectively, led by Drs. Mingder Yang and Irina Kusmartseva for assisting with organ procurement and processing. This work was supported by the National Institutes of Health (NIH) R01-DK123292 and R01-DK131059, the DFG Collaborative Research Centre/Transregio 127, the DFG — International Research Training Group 2251, JRDF grant 2- SRA-2019-696-S-B, the Paul Langerhans Institute Dresden (PLID) of the Helmholtz Zentrum München at the University Clinic Carl Gustav Carus of Technische Universität Dresden, and the German Ministry for Education and Research to the German Centre for Diabetes Research. We also acknowledge the support of the nPOD (RRID:SCR_014641), a collaborative T1D research project sponsored by JDRF (nPOD: 5-SRA-2018-557-Q-R), The Leona M. and Harry B. Helmsley Charitable Trust (grant 2018PG-T1D053, 2018PG-T1D060, and 2015PG-T1D052), and by the Diabetes Research Institute Foundation. The content and views expressed are the responsibility of the authors and do not necessarily reflect the official view of nPOD. Organ Procurement Organizations (OPO) partnering with nPOD to provide research resources are listed at http://www.jdrfnpod.org//for-partners/npod-partners. Finally, we wholeheartedly thank the donors and their families for their invaluable contribution to science.

## Author contributions

Conceptualization: S.S., M.A.A. Methodology: D.M.D., A.K.M.A.

Investigation: D.M.D., A.K.M.A., D.T.B., C.M.C., C.T.M.

Writing – Original Draft: D.M.D., A.K.M.A., D.T.B.

Writing – Review & Editing: D.M.D, A.K.M.A., C.M.C., C.T.M., A.L.P., M.D.W., M.A.B., C.H.W, S.S, M.A.A.

Funding Acquisition: E.A.P., S.S., M.A.A. Supervision: M.A.B., C.H.W, S.S., M.A.A.

## STAR METHODS

### KEY RESOURCES TABLE

#### LEAD CONTACT AND MATERIALS AVAILABILITY

This study did not generate new unique reagents or original code. Further information and requests for resources and reagents should be directed to and will be fulfilled by the Lead Contact, Mark Atkinson.

#### EXPERIMENTAL MODEL AND SUBJECT DETAILS

All procedures were performed according to the established standard operating procedures of the nPOD/OPPC and approved by the University of Florida Institutional Review Board (IRB201600029) and the United Network for Organ Sharing (UNOS) according to federal guidelines, with informed consent obtained from each donor’s legal representative. Pancreas tissue from human donors with or without T1D was procured from the nPOD program at the University of Florida (RRID: SCR_014641, https://www.jdrfnpod.org). For each donor, a medical chart review was performed, and C-peptide was measured, with T1D diagnosed according to the guidelines established by the American Diabetes Association (ADA). Demographic data, hospitalization duration, and organ transport time were obtained from hospital records. Donor pancreata were recovered, placed in transport media on ice, and shipped via organ courier to the University of Florida. The tissue was processed by a licensed pathology assistant. Detailed donor information is listed in Supplemental Table 1.

### METHOD DETAILS

#### Living pancreas tissue slice preparation

Fresh, living tissue slices from organ donor pancreata were prepared as previously described^29^. Briefly, cross-sections from the PH, PB, and PT regions of 27 human pancreata (Supplemental Table 1) were freshly obtained from nPOD (https://npod.org) and placed into separate 100 mm dishes containing ECS (125 mM NaCl, 26 mM NaHCO_3_, 10 mM HEPES, 3 mM glucose, 2.5 mM KCl, 2 mM CaCl_2_, 1.25 mM NaH_2_PO_4_, 1 mM MgCl_2_, pH 7.4) supplemented with aprotinin (25 KIU/mL). Small tissue pieces approximately 0.5 cm^3^ in size were cut from the larger cross sections, and any remaining connective, fibrotic, or adipose tissue was removed before embedding the pieces in low-melting point agarose (3.8%). The agarose solidified at 4- 8°C and the resulting tissue blocks were mounted onto the specimen holder of a semi-automated vibratome. The tissue blocks were sliced at a thickness of 120 µm, speed of 0.1-0.2 mm/s, amplitude of 1 mm, and angle of 15 degrees. Slices were collected in separate dishes containing ECS with aprotinin. After slices from all regions were prepared, they were transferred to dishes containing Low Glucose DMEM (Dulbecco’s Modified Eagles Medium) supplemented with 10% FBS, antibiotic-antimycotic solution (1:100 dilution), and 25 KIU/mL aprotinin, and placed in an incubator set at 28°C with 5% CO_2_ for overnight resting.

#### Dynamic perifusion assay for insulin and glucagon secretion measurement

Tissue slices were placed in 3mM glucose containing DMEM media supplemented with 3.2 g/L NaHCO_3_, 1.11 g/L HEPES, 0.11 g/L sodium pyruvate, 4 mM L-glutamine, pH 7.4, 0.1% BSA and 25 KIU/mL aprotinin for 2 hours, gently rocking. After the 2-hour resting period, slices from the PH, PB, and PT regions were loaded into separate perifusion chambers (4 slices/chamber/region) and connected to a perifusion system. Tissue slices were perifused at a flow rate of 100 µL/minute at 37°C, and the perifusate was collected in 96-well plates at 2- minute intervals. The slices were first flushed with media containing 3 mM glucose for 60 minutes to wash out accumulated hormones and enzymes, and then subjected to a series of glucose challenges – 10 minutes of media with 1 mM glucose, 20 minutes with 5.5 mM glucose, 20 minutes with 1 mM glucose, 20 minutes with 11.1 mM glucose, 20 minutes with 1 mM glucose, 10 minutes with 11.1 mM glucose and 30 mM KCl, and 10 minutes with 1 mM glucose. Perifusate samples were stored at -20°C until measurement of insulin and glucagon with commercially available ELISA kits.

After perifusion, slices were removed from the chambers and fixed in 4% paraformaldehyde for 30 minutes, then washed in PBS for 5 minutes. The slices were then transferred into 5 mL tubes filled with PBS with azide and stored at 4°C until shipment to the Paul Langerhans Institute Dresden for 3D morphometric analysis.

#### Assessment of pancreatic amylase, lipase, and trypsinogen release

Five slices from the PH, PB, and PT regions were incubated in 3 mM glucose containing DMEM media, supplemented with aprotinin on an orbital shaker (15-20 rpm) at 37°C for one hour. After the one-hour resting period, slices were transferred to a 24-well plate with 500 µL of fresh media in each well. Basal secretion samples were collected after a 30-minute incubation at 37°C on an orbital shaker. Slices were then stimulated with 10 µM carbachol for 30 minutes at 37°C on an orbital shaker. After the stimulation step the supernatants were collected, and the slices were placed into 500 µL 3 mM DMEM with 3% Triton X-100 for the assessment of total slice enzyme content. All samples were stored at -20°C until amylase, lipase and trypsinogen measurement using commercially available ELISA and RIA kits.

#### Immunofluorescent staining and whole slice 3D imaging

Whole slices were immersed in a blocking solution (GSDB) containing 0.3% Triton X-100 (1% goat serum, 0.3% Triton X-100, 900 mM NaCl, and 40 mM sodium phosphate buffer in deionized water with a pH of 7.4) and stained using primary antibodies targeting insulin (1:10 dilution) and glucagon (1:2000 dilution) for 24-48 hours at 4°C with gentle shaking. After primary antibody incubation, the slices were washed three times for at least 30 minutes each with PBS. Subsequently, the slices were incubated with secondary antibodies: AlexaFluor 488 goat anti-guinea pig (1:200 dilution) for insulin, AlexaFluor 633 goat anti-mouse (1:200 dilution) for glucagon, and DAPI (2.5mg/ml) overnight at 4°C with gentle shaking. Following secondary antibody and DAPI incubation, the slices were washed three times for at least 30 minutes. Slices were then stored in PBS at 4°C in the dark until imaging.

Stained slices were mounted underneath a glass coverslip, submerged in PBS (pH 7.4), and imaged in their entirety using an upright laser scanning confocal microscope (TCS SP8 SMD, Leica) equipped with a water-immersion 20x high NA objective lens and a motorized stage. Single tiles were imaged as previously described^29^, at a resolution of 0.75×0.75 mm, with a tile overlap of 15% and a stack separation of 2.5 mm. DAPI and fluorescent antibodies were excited at 405nm, 488nm and 633nm, and detected at 405-510nm (for DAPI), 490-560nm (insulin), and 638-755nm (glucagon), respectively.

#### 3D#morphometric analysis of insulin and glucagon volumes

Tilescans were stitched using the built-in stitching function of the Las X Life Science Microscope Software Platform. Data quantification was semi-automatically conducted using a customized FIJI toolset, as previously described^26^. Briefly, total slice area and individual endocrine objects (defined as those positive for at least one hormone stain) were manually contoured on a maximum intensity projection, and regions of interest (ROIs) were saved separately. For individual endocrine particles, regions were cropped from the stitched tilescan, followed by subtraction of the 488 channel from the 546 channel and removal of the 546 channel from the 633 channel to account for channel bleed. Images were then split by color, contrast-enhanced (CLAHE), involving median filtering (3×3×1) and conversion to binary (IJ_IsoData).

Further processing included hole filling in 3D, size opening (limited to 500 μm^3^), and volume analysis (Euler connectivity: C26) using MorphoLibJ^45^. The total slice volume was calculated by adding each channel of the original tilescan and cropping it using the previously contoured total slice ROI. Additional steps included median filtering (3×3×1), mask generation for nearby points (within a distance of 10.0 μm, threshold of 12.75), removal of dark and bright outliers, hole filling, 3D volume closing (cube, 3×3×1), and volume analysis for endocrine objects using MorphoLibJ^45^.

### QUANTIFICATION AND STATISTICAL ANALYSIS

#### Statistical analyses and Calculations

No specific statistical methods were used to predetermine sample size due to no ability to predict sample availability. All results are presented as mean ± SEM. The significance of the differences between groups was analyzed as described in the figure legends. P values < 0.05 were considered statistically significant. All statistical analyses were performed using GraphPad Prism Software version 9.

### DATA AND CODE AVAILABILITY

This study did not generate any unique datasets or code.

## Supplemental Information titles and legends

**Table S1. Characteristics of ND, AAb+, and T1D organ donors with secretion data collected from head, body, and tail pancreas regions, related to STAR Methods**

**Figure S1. Characteristics of organ donors used for functional studies, Related to Table S1**

Donor age (A), body mass index (B), hemoglobin A1c (C) and serum C-peptide (D) levels. N=15 ND, 7 1AAb+, and 5 T1D donors

Dots represent individual donors. Data are represented as mean ± SEM. Significance was assessed using two-way ANOVA of log-transformed data. ∗p < 0.05; ∗∗p < 0.01; ∗∗∗p < 0.001; ∗∗∗∗p < 0.0001; ns, not significant.

**Figure S2. Insulin and glucagon secretion in AAb+ and recent-onset T1D, Related to Figure 2**

A-B. Insulin secretion traces from slices of ND (blue), 1AAb+ (orange) and recent-onset T1D (red) donors (mean of pancreas head, body and tail) shown as absolute secreted amounts (A) and stimulation index calculated as fold of baseline at 1G (B).

C-E. Quantification of insulin responses to 5.5G (C), 11.1G (D), and KCl (E) stimulation.

F-G. Glucagon secretion traces from slices of ND (blue), 1AAb+ (orange) and recent-onset T1D (red) donors (mean of pancreas head, body and tail) shown as absolute secreted amounts (F) and stimulation index calculated as fold of baseline at 5.5G (G).

H-K. Quantification of glucagon responses to 1G (H), 11.1G (I), 1G (J) and KCl (K) stimulation. N=15 ND, 7 1AAb+, and 5 T1D donors, with 4 slices/region/donor

Dots represent individual donors. Data are represented as mean ± SEM. Significance was assessed using two-way ANOVA of log-transformed data. ∗p < 0.05; ∗∗p < 0.01; ∗∗∗p < 0.001; ∗∗∗∗p < 0.0001; ns, not significant.

**Figure S3. Insulin and glucagon secretion across all pancreas regions in AAb+ and recent-onset T1D, Related to Figure 2**

A-B. Insulin secretion traces from slices of the pancreas head, body, and tail of 1AAb+ donors shown as absolute secreted amounts (A) and stimulation index calculated as fold of baseline at 1G (B).

C-E. Quantification of insulin responses to 5.5G (C), 11.1G (D), and KCl (E) stimulation.

F-G. Insulin secretion traces from slices of the pancreas head, body and tail of T1D donors shown as absolute secreted amounts (F) and stimulation index calculated as fold of baseline at 1G (G).

H-J. Quantification of insulin responses to 5.5G (H), 11.1G (I), and KCl (J) stimulation.

K-L. Glucagon secretion traces from slices of the pancreas head, body and tail of 1AAb+ donors shown as absolute secreted amounts (K) and stimulation index calculated as fold of baseline at 5.5G (L).

M-P. Quantification of glucagon responses to 1G (M), 11.1G (N), 1G (O) and KCl (P) stimulation.

Q-R. Glucagon secretion traces from slices of the pancreas head, body and tail of T1D donors shown as absolute secreted amounts (Q) and stimulation index calculated as fold of baseline at 5.5G (R).

S-V. Quantification of glucagon responses to 1G (S), 11.1G (T), 1G (U) and KCl (V) stimulation. N=15 ND, 7 1AAb+, and 5 T1D donors, with 4 slices/region/donor

Dots represent individual donors. Data are represented as mean ± SEM. Significance was assessed using two-way ANOVA of log-transformed data. ∗p < 0.05; ∗∗p < 0.01; ∗∗∗p < 0.001; ∗∗∗∗p < 0.0001; ns, not significant.

