## Supplementary material for "Impaired islet function with normal exocrine enzyme secretion is consistent across the head, body, and tail pancreas regions in type 1 diabetes": Key Resources Table

| REAGENT or RESOURCE | SOURCE | IDENTIFIER |
| --- | --- | --- |
| Antibodies | | |
| Polyclonal Guinea Pig Anti-Insulin | Agilent - Dako | Cat# IR00261-2  RRID: AB_2800361 |
| Monoclonal Mouse Anti-Glucagon | Sigma-Aldrich | Cat# G2654  RRID: AB_259852 |
| AlexaFluor® 488 Goat Anti-Guinea Pig | Thermo Fisher Scientific - Invitrogen | Cat# A-11073  RRID: AB_2534117 |
| AlexaFluor® 633 Goat Anti-Mouse | Thermo Fisher Scientific - Invitrogen | Cat# A-21050  RRID: AB_2535718 |
| Bacterial and virus strains | | |
| Biological samples |  |  |
| Human Pancreas Tissue | nPOD; https://npod.org/ | RRID: SCR_014641 |
| nPOD-6540 |  | SAMN25652251 |
| nPOD-6543 |  | SAMN25652254 |
| nPOD-6544 |  | SAMN25652255 |
| nPOD-6546 |  | SAMN25652257 |
| nPOD-6547 |  | SAMN25652258 |
| nPOD-6548 |  | SAMN25652259 |
| nPOD-6550 |  | SAMN25652261 |
| nPOD-6551 |  | SAMN25652262 |
| nPOD-6552 |  | SAMN30386842 |
| nPOD-6553 |  | SAMN30386843 |
| nPOD-6556 |  | SAMN30386845 |
| nPOD-6558 |  | SAMN30386847 |
| nPOD-6559 |  | SAMN30386848 |
| nPOD-6560 |  | SAMN30386849 |
| nPOD-6562 |  | SAMN30386850 |
| nPOD-6563 |  | SAMN30386851 |
| nPOD-6569 |  | SAMN38117299 |
| nPOD-6571 |  | SAMN33284289 |
| nPOD-6573 |  | SAMN33284291 |
| nPOD-6575 |  | SAMN33284293 |
| nPOD-6578 |  | SAMN33284295 |
| nPOD-6579 |  | SAMN33284296 |
| nPOD-6582 |  | SAMN38117302 |
| nPOD-6583 |  | SAMN38117303 |
| nPOD-6584 |  | SAMN38117304 |
| nPOD-6586 |  | SAMN38117306 |
| nPOD-CV14 |  | SAMN38117313 |
| Chemicals, peptides, and recombinant proteins | | |
| Aprotinin | MilliporeSigma | Cat# A6106 |
| Low Melting Point Agarose | MilliporeSigma | Cat# A0701 |
| Low Glucose DMEM | MilliporeSigma | Cat# D6046 |
| Fetal Bovine Serum | Cytiva | Cat# SH30071.03IH |
| Antibiotic-Antimycotic Solution | Corning | Cat# 30-004-CI |
| DMEM (Powder) | Corning | Cat# MT9013PB |
| Sodium Pyruvate | MilliporeSigma | Cat# P2256 |
| HEPES | MilliporeSigma | Cat# H4034 |
| L-Glutamine | Thermo Fisher | Cat# 25030081 |
| Bovine Serum Albumin | MilliporeSigma | Cat# A6003 |
| Carbamylcholine chloride | MilliporeSigma | Cat# C4382 |
| Triton X-100 | MilliporeSigma | Cat# 648462 |
| 4% Paraformaldehyde | Thermo Fisher Scientific | Cat# J19943.K2 |
| PBS with Azide | Santa Cruz Biotechnology | Cat# SC-296028 |
| DAPI | Sigma-Aldrich | Cat# D9542 |
| Critical commercial assays | | |
| Insulin ELISA Kit | Mercodia | Cat# 10-1113-10 |
| Ultrasensitive Insulin ELISA | Mercodia | Cat# 10-1132-01 |
| Glucagon ELISA Kit | Crystal Chem | Cat# 81520 |
| Human Pancreatic Amylase ELISA Kit | Abcam | Cat# ab137969 |
| Human Pancreatic Lipase ELISA Kit | Novus Biologicals | Cat# NB016815 |
| Trypsin RIA | ALPCO | Cat# 72-OCFE07-TRYPS |
| Deposited data | | |
| Experimental models: Cell lines | | |
| Experimental models: Organisms/strains | | |
| Oligonucleotides | | |
| Recombinant DNA | | |
| Software and algorithms | | |
| Prism 9.4.0 | GraphPad | RRID: SCR_002798 |
| ImageJ | Schneider et al., 2012 | https://imagej.nih.gov/ij/ |
| MorphoLibJ | Legland et al., 2016 | https://imagej.net/plugins/morpholibj |
| FIJI | Schindelin et al., 2012 | https://imagej.net/software/fiji/ |
| Las X Life Science Microscope Software Platform | Leica | https://www.leica-microsystems.com/products/microscope-software/p/leica-las-x-ls/ |
| Other | | |
| Perifusion System | Biorep Technologies | Cat#PERI4-115 |
| Semiautomatic Vibratome VT1200S | Leica | Cat#14048142065 |
| Perifusion Chamber | Warner Instruments | Cat#64-0223 |
| Perifusion Platform | Warner Instruments | Cat#64-0281 [P-5] |
| Laser Scanning Confocal Microscope | Leica | Model: TCS SP8 SMD |
