## Supplemental Materials for "Impaired islet function with normal exocrine enzyme secretion is consistent across the head, body, and tail pancreas regions in type 1 diabetes"

**Table S1. Characteristics of ND, AAb+, and T1D organ donors with secretion data collected from head, body, and tail pancreas regions.**

| Donor no. | nPOD ID | Donor Type | Autoantibody status | Age (years) | Diabetes Duration (months) | Sex | C-peptide (ng/ml) | HbA1c (%) | BMI |
|---|---|---|---|---|---|---|---|---|---|
| **1** | 6540 | ND | Negative | 6.86 | N/A | F | 2.03 | 5.8 | 18.3 |
| **2** | 6543 | ND | Negative | 3.50 | N/A | M | 1.45 | 5.3 | 17.4 |
| **3** | 6544 | ND | Negative | 12.45 | N/A | M | 8.66 | 5.2 | 23.4 |
| **4** | 6546 | ND | Negative | 22.29 | N/A | M | 11.00 | 5.6 | 23.7 |
| **5** | 6547 | ND | Negative | 6.64 | N/A | F | 3.89 | 5.5 | 24.8 |
| **6** | 6548 | ND | Negative | 20.24 | N/A | M | 4.04 | 5.7 | 23.8 |
| **7** | 6552 | ND | Negative | 33.87 | N/A | F | 1.80 | 5.6 | 21.9 |
| **8** | 6556 | ND | Negative | 34.60 | N/A | M | 11.43 | 5.5 | 30.8 |
| **9** | 6559 | ND | Negative | 23.46 | N/A | F | 7.02 | 5.3 | 26.3 |
| **10** | 6560 | ND | Negative | 34.36 | N/A | F | 10.36 | 5.9 | 26.6 |
| **11** | CV14 | ND | Negative | 9.00 | N/A | M | 4.40 | 5.7 | 13.7 |
| **12** | 6571 | ND | Negative | 28.10 | N/A | M | 4.57 | 6.1 | 29.7 |
| **13** | 6583 | ND | Negative | 30.54 | N/A | M | 10.55 | 5.0 | 24.8 |
| **14** | 6584 | ND | Negative | 22.53 | N/A | M | 6.02 | 5.3 | 21.1 |
| **15** | 6586 | ND | Negative | 13.21 | N/A | M | 6.83 | 5.1 | 27.4 |
| **16** | 6553 | AAb+ | mIAA+ | 12.34 | N/A | F | 4.62 | 5.7 | 24.8 |
| **17** | 6558 | AAb+ | GAD+ | 21.69 | N/A | F | 8.03 | 4.4 | 27.8 |
| **18** | 6562 | AAb+ | GAD+ | 29.77 | N/A | F | 13.19 | 5.6 | 18.2 |
| **19** | 6569 | AAb+ | GAD+ | 20.00 | N/A | F | 2.44 | 6.0 | 24.2 |
| **20** | 6573 | AAb+ | GAD+ | 24.48 | N/A | F | 1.85 | 5.1 | 35.5 |
| **21** | 6575 | AAb+ | GAD+ | 23.53 | N/A | M | 5.21 | 5.5 | 26.9 |
| **22** | 6582 | AAb+ | GAD+ | 22.38 | N/A | F | 2.67 | 5.6 | 23.0 |
| **23** | 6550 | T1D | GAD+ ZnT8+ | 25.06 | 0 | M | <0.02 | 14.0 | 16.4 |
| **24** | 6551 | T1D | GAD+ IA-2+ mIAA+ ZnT8+ | 20.70 | 7 | M | 0.11 | 6.4 | 23.1 |
| **25** | 6563 | T1D | IA-2+ | 14.60 | 0 | F | 1.04 | 9.6 | 25.5 |
| **26** | 6578 | T1D | IA2+ ZnT8+ | 11.95 | 0 | F | 0.35 | 13.6 | 22.2 |
| **27** | 6579 | T1D | GAD+ mIAA+ | 13.91 | 14 | F | 0.31 | 15.0 | 18.4 |

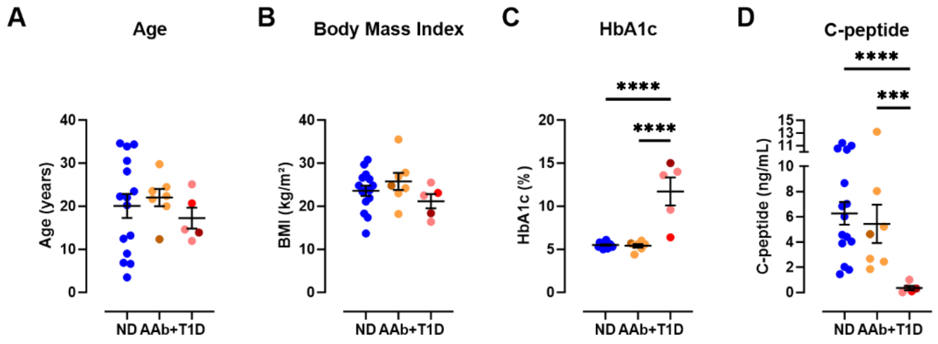


**Figure S1. Characteristics of organ donors used for functional studies, Related to Table S1**

Donor age (A), body mass index (B), hemoglobin A1c (C) and serum C-peptide (D) levels.

N=15 ND, 7 1AAb+ and 5 T1D donors

Dots represent individual donors. Data are represented as mean ± SEM. Significance was assessed using two-way ANOVA of log-transformed data. ∗$p < 0.05$; ∗∗$p < 0.01$; ∗∗∗$p < 0.001$; ∗∗∗∗$p < 0.0001$; ns, not significant.

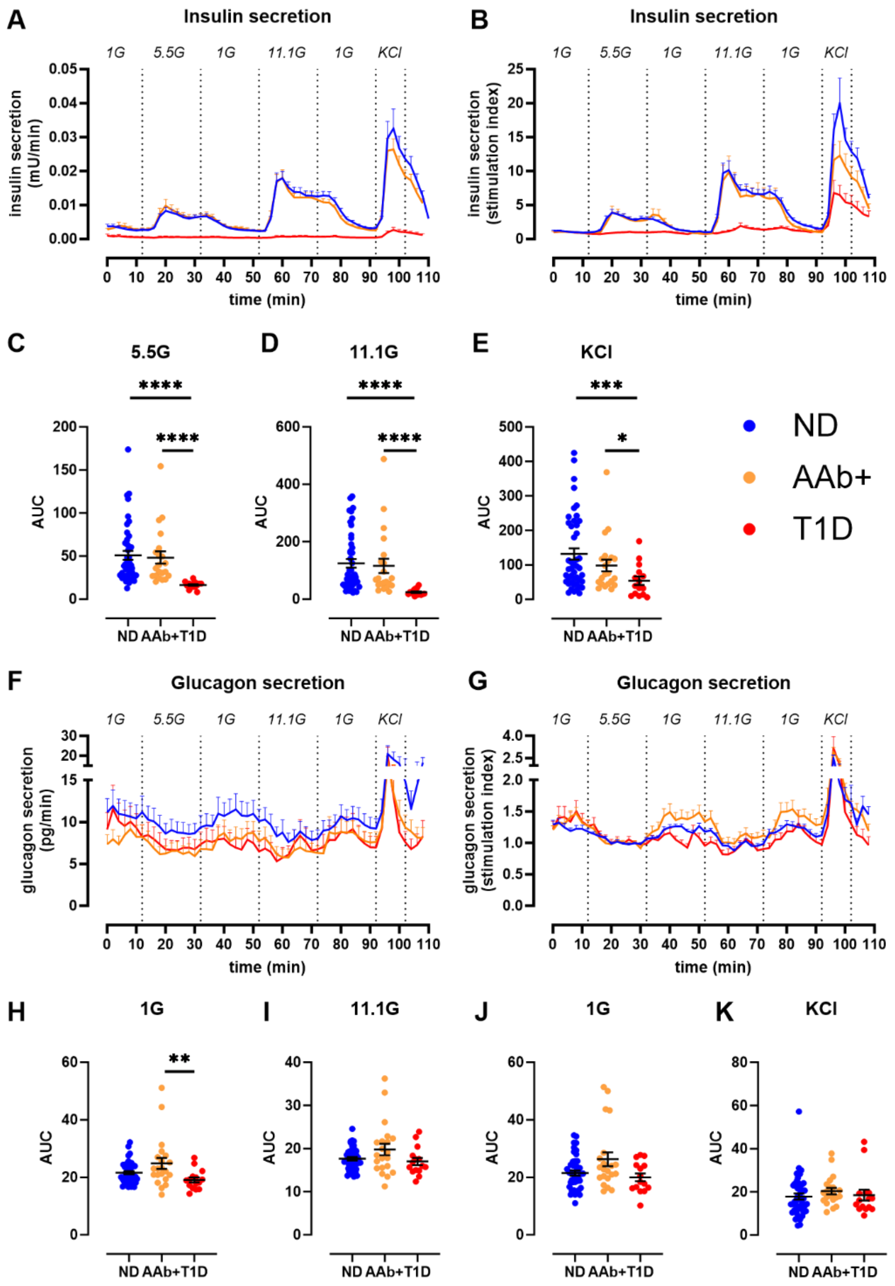


**Figure S2. Insulin and glucagon secretion in AAb+ and recent-onset T1D, Related to Figure 2**

A-B. Insulin secretion traces from slices of ND (blue), 1AAb+ (orange) and recent-onset T1D (red) donors (mean of pancreas head, body and tail) shown as absolute secreted amounts (A) and stimulation index calculated as fold of baseline at 1G (B).

C-E. Quantification of insulin responses to 5.5G (C), 11.1G (D) and KCl (E) stimulation.

F-G. Glucagon secretion traces from slices of ND (blue), 1AAb+ (orange) and recent-onset T1D (red) donors (mean of pancreas head, body and tail) shown as absolute secreted amounts (F) and stimulation index calculated as fold of baseline at 5.5G (G).

H-K. Quantification of glucagon responses to 1G (H), 11.1G (I), 1G (J) and KCl (K) stimulation.

N=15 ND, 7 1AAb+ and 5 T1D donors, with 4 slices/region/donor

Dots represent individual donors. Data are represented as mean ± SEM. Significance was assessed using two-way ANOVA of log-transformed data. $*p < 0.05$; $**p < 0.01$; $***p < 0.001$; $****p < 0.0001$; ns, not significant.

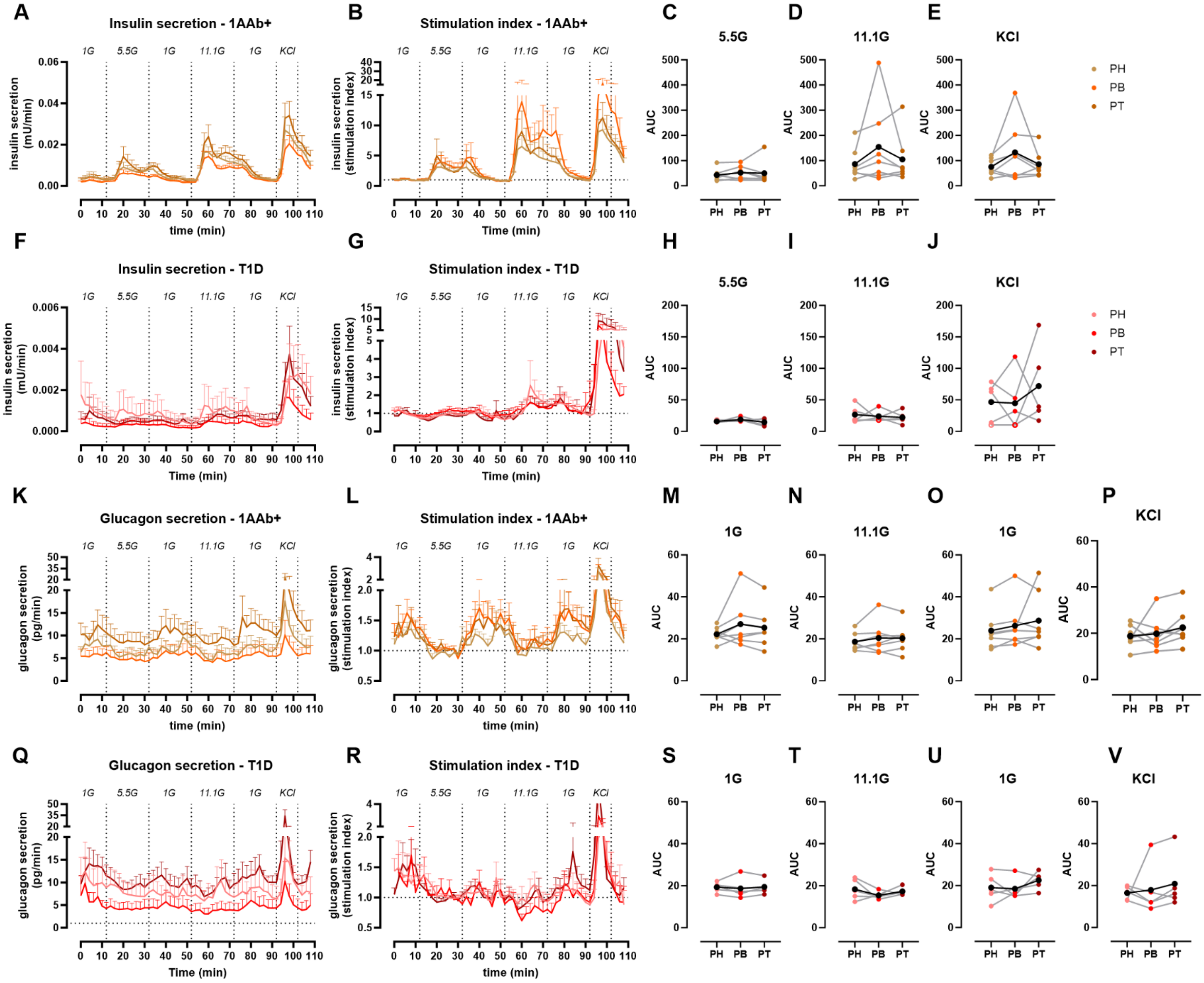


**Figure S3. Insulin and glucagon secretion across all pancreas regions in AAb+ and recent-onset T1D, Related to Figure 2**

A-B. Insulin secretion traces from slices of the pancreas head, body and tail of 1AAb+ donors shown as absolute secreted amounts (A) and stimulation index calculated as fold of baseline at 1G (B).

C-E. Quantification of insulin responses to 5.5G (C), 11.1G (D) and KCl (E) stimulation.

F-G. Insulin secretion traces from slices of the pancreas head, body and tail of T1D donors shown as absolute secreted amounts (F) and stimulation index calculated as fold of baseline at 1G (G).

H-J. Quantification of insulin responses to 5.5G (H), 11.1G (I) and KCl (J) stimulation.

K-L. Glucagon secretion traces from slices of the pancreas head, body and tail of 1AAb+ donors shown as absolute secreted amounts (K) and stimulation index calculated as fold of baseline at 5.5G (L).

M-P. Quantification of glucagon responses to 1G (M), 11.1G (N), 1G (O) and KCl (P) stimulation.

Q-R. Glucagon secretion traces from slices of the pancreas head, body and tail of T1D donors shown as absolute secreted amounts (Q) and stimulation index calculated as fold of baseline at 5.5G (R).

S-V. Quantification of glucagon responses to 1G (S), 11.1G (T), 1G (U) and KCl (V) stimulation.

N=15 ND, 7 1AAb+ and 5 T1D donors, with 4 slices/region/donor

Dots represent individual donors. Data are represented as mean ± SEM. Significance was assessed using two-way ANOVA of log-transformed data. $*p < 0.05$; $**p < 0.01$; $***p < 0.001$; $****p < 0.0001$; ns, not significant.
